# Recent Cis-Trans Coevolution Driven by the Emergence of A Novel Gene in Drosophila

**DOI:** 10.1101/045385

**Authors:** Benjamin H Krinsky, Robert K. Arthur, Kevin P. White, Manyuan Long

**Affiliations:** Committee on Evolutionary Biology, University of Chicago, Chicago, Illinois, United States of America; Department of Ecology and Evolution, University of Chicago, Chicago, Illinois, United States of America; Institute for Genomics and Systems Biology, Department of Human Genetics, University of Chicago and Argonne National Laboratory, Chicago, Illinois, United States of America; Current Address: Federation of American Societies for Experimental Biology (FASEB), Bethesda, Maryland, United States of America

## Abstract

Young, or newly evolved, genes arise ubiquitously across the tree of life, and can rapidly acquire novel functions that influence a diverse array of biological processes^1^. Previous work identified a young regulatory gene in *Drosophila*, *Zeus*, which diverged rapidly from its parent *Caf40* and took on roles in the male reproductive system. This neofunctionalization was accompanied by differential binding of the Zeus protein to loci throughout the *Drosophila melanogaster* genome^2^. However, the way in which new DNA-binding proteins acquire and coevolve with their targets in the genome is not understood. Here, by comparing Zeus ChIP-seq data from *D. melanogaster* and *D. simulans* to the ancestral Caf40 binding events from *D. yakuba*, a species that diverged before the duplication event, we find a dynamic pattern in which Zeus binding rapidly co-evolved with a previously unknown DNA motif under the influence of positive selection. Interestingly, while both copies of Zeus acquired targets at male-biased and testis-specific genes, *D. melanogaster* and *D. simulans* proteins have specialized binding on different chromosomes, a pattern echoed in the evolution of the associated motif. Our results suggest that evolution of young regulatory genes can be coupled to substantial rewiring of the transcriptional networks into which they integrate, even over short evolutionary timescales. Our results thus uncover dynamic, genome-wide evolutionary processes associated with new genes.

## Introduction

The origin of new genes can lead to the evolution of new and crucial functions in myriad biological processes, including gene regulation^1,3,4^. Regulatory and other putative functional elements can now be investigated on a genome-wide scale in order to systematically characterize networks of gene-gene interactions^5^, and to compare patterns of conservation and divergence of gene regulation in multiple, closely related species^6–8^. However, most of the comparisons to date have focused on conserved factors with well-characterized molecular functions (Ni et al. 2012, Schmidt et al. 2010). Thus, there exists a unique opportunity to apply these new approaches to investigate the evolution of new regulatory genes and as well as their effects on bound regulatory elements, and therefore explore how newly arisen loci might evolve altered or gene-gene interactions across the genome.

The gene *Zeus* (CG9573, also known as Rcd-1r) is a testis-specific young gene that arose via a retroposition event approximately 5 million years ago in the lineage leading to *Drosophila melanogaster* and its closest relatives. *Zeus* subsequently underwent a very rapid period of molecular evolution^9, 10^. Functional analyses suggest that *Zeus* evolved specific roles in the development and function of *Drosophila* sperm and testis^2^. This evolution in *Zeus*’ function coincided with changes in its expression and patterns of histone modification at the Zeus locus^11^.

In contrast, its parental gene *Caf40* (CG14213, also known as Rcd-1), conserved across eukaryotes, is ubiquitously expressed and is essential for viability in *Drosophila melanogaster*^2^. On the molecular level, it had been previously inferred that Caf40 has nucleic acid binding properties, and thus might act as a regulator through its interactions with genomic DNA^12^. By performing chromatin immunoprecipitation followed by microarray analysis (ChIP-chip), it was subsequently discovered that both Zeus and Caf40 from *D. melanogaster* bind to several hundred sites throughout the genome, and that Zeus has acquired a number of novel regulatory targets in the genome, consistent with neofunctionalization following duplication^2^.

The *Zeus* locus arose after the divergence of the lineages leading to *D. melanogaster* and *D. yakuba*, but prior to the divergence of *D. melanogaster* and one of its closest sister species, *D. simulans*. The Zeus protein subsequently acquired a large number of species-specific substitutions along these two lineages^2^ (Fig. 1A). To understand patterns of lineage specific regulatory evolution of Zeus, as well as its initial divergence from the ancestral state of Caf40, we have characterized the genome-wide binding profiles of Zeus from *D. melanogaster* and its sister species *D. simulans*, as well as Caf40 from *D. yakuba* using chromatin immunoprecipitation followed by high-throughput sequencing (ChIP-seq). We selected *D. yakuba* (pre-duplication) Caf40 as the best proxy from which to infer ancestral Caf40 binding, because the *D. yakuba* and *D. melanogaster* proteins differ in only 4 positions.

**Figure 1.**
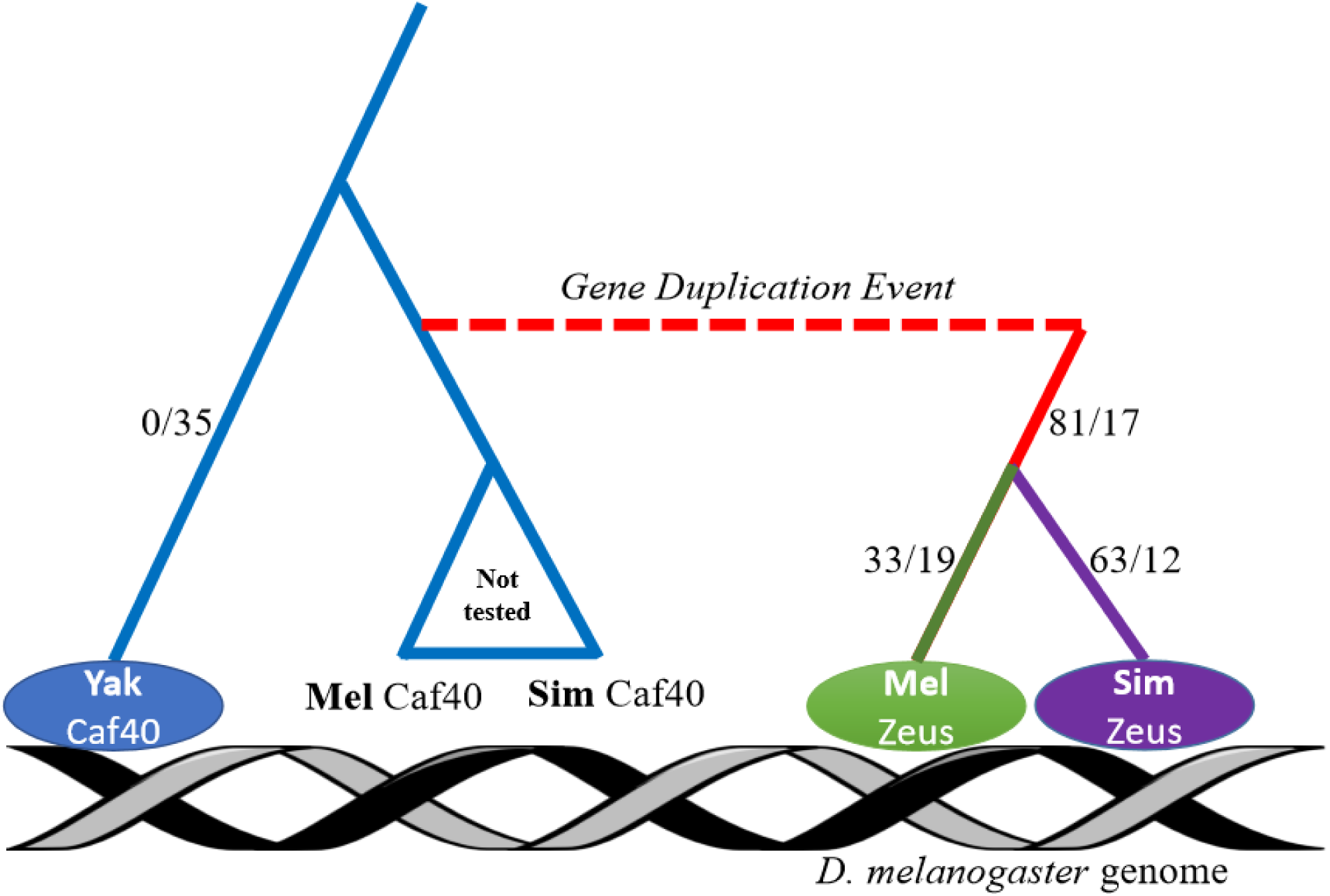
Design and results of ChIP-seq experiments.Depiction of Zeus/Caf-40 phylogeny, with experimental design. Zeus originated from a gene duplication event 4-6 million years ago, before the split of *D. melanogaster* and *D. yakuba*. We sampled two copies of Zeus (*D. melanogaster* and *D. simulans*) as well as a single copy of Caf-40 from *D. yakuba*, which represents the ancestral, pre-duplication state of the protein. All three proteins were introduced into the *D. melanogaster* genome with 3x FLAG tags attached, in order to eliminate problems with variable antibody affinity. Numbers indicate the volume of nonsynonymous (before the slash) and synonymous (after the slash) changes.

## Results

We engineered transgenic lines of *D. melanogaster* (w1118) containing FLAG-tagged *D. melanogaster* Zeus, *D. simulans* Zeus and *D. yakuba* Caf40, and performed ChIP-seq on each of these lines (Fig. 1). This allows us to directly compare binding properties of the three proteins in a common genome. We term these three proteins Dmel Zeus, Dsim Zeus, and Dyak Caf, respectively. We obtained reproducible ChIP-seq signals between replicates (Supplementary Fig. 1; Supplementary Table 1), and observed Zeus binding affinity correlated with temporal patterns of expression in the testis (Supplementary Fig. 2). We observed strong enrichment of ChIP signal primarily at the transcription start site and within the exons of bound genes (Supplementary Fig. 3), refining previously hypothesized Zeus and Caf40 binding preferences^2^. To assess the potential differences in binding among the three proteins, we calculated signal enrichment for each gene based on the enrichment of reads within 700bp of the transcription start site (TSS). Principal component analysis on the gene-by-gene signal revealed that replicates corresponding to each protein (Dmel Zeus, Dsim Zeus,Dyak Caf40) formed distinct clusters (Fig. 2A), demonstrating significant differences in binding preferences between proteins. We computed the pairwise Euclidean distance between proteins’ read counts, which showed that Dsim Zeus sites were more highly diverged from Dyak Caf40 than Dmel Zeus sites (Supplementary Fig. 4; p<.001), consistent with the reported pattern of protein-coding sequence divergence (Fig. 1).

**Figure 2.**
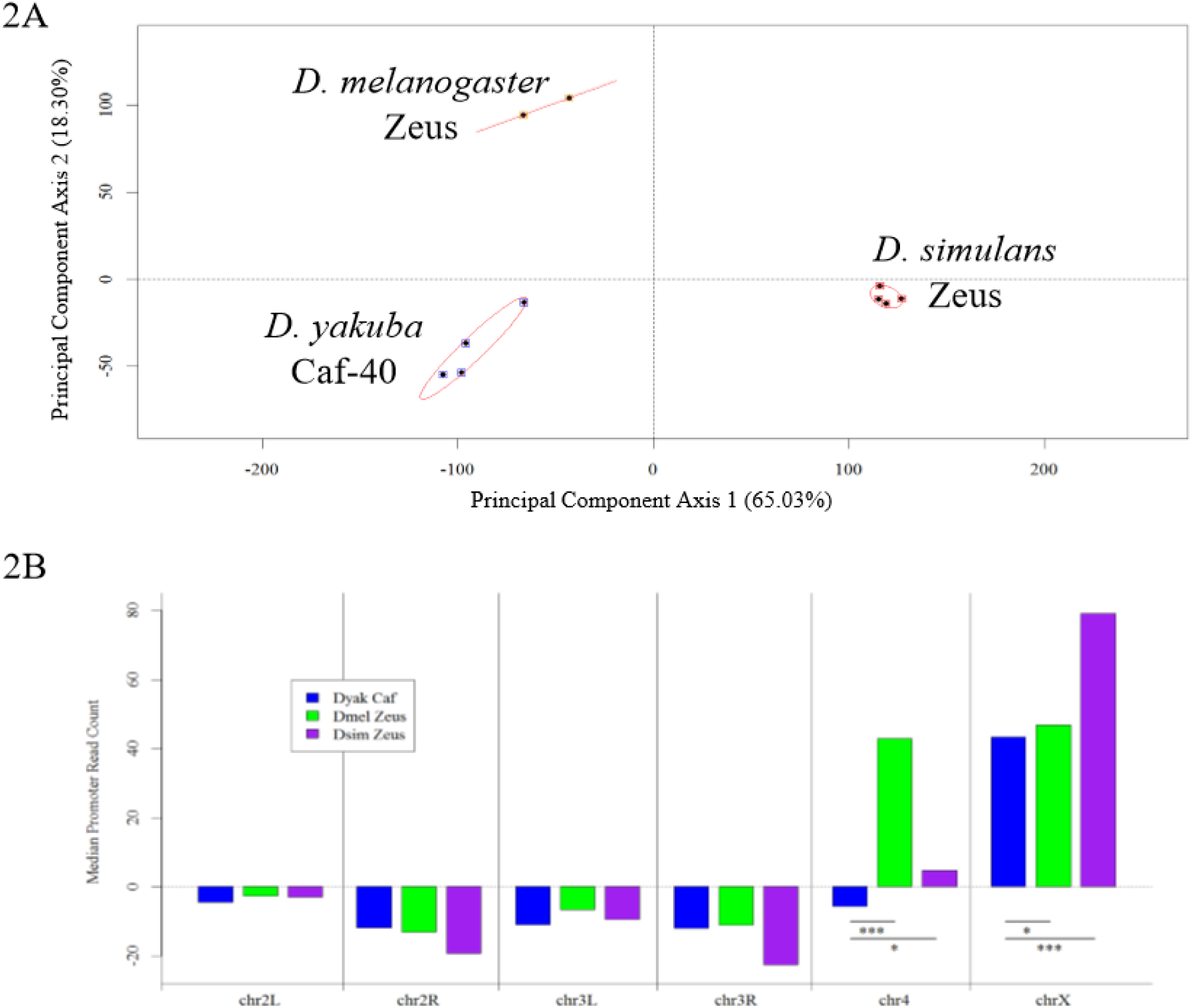
The Evolution of Zeus Binding Affinity in Trans. A) A graph of the first two principal components of ChIP-seq read counts revealed reproducible clustering of replicates of the same protein, while different proteins showed differentiation. B) A bar plot showing the median normalized read counts over TSSs for each chromosome, indicating differences in chromosome-level affinity of the three proteins. Both copies of Zeus show increased affinity relative to Caf-40 on chromosomes X and 4, albeit to different degrees.

Based on previous ChIP-chip results showing that Zeus preferentially binds the X-chromosome, and because of the known roles of Zeus in regulating sex-specific functions (which are enriched on the X-chromosome), we compared the chromosomal distribution of reads^13^. We found ChIP-seq read enrichment for all three proteins on the X-chromosome relative to the autosomes (Fig. 2B), but both Zeus orthologs showed significantly higher X vs. autosome signal enrichment compared to Dyak Caf40 (Permutation test: p<.05). Dsim Zeus showed particularly strong X-chromosome enrichment (Permutation test: p<.001).

Both Zeus proteins also exhibited a bias for the 4^th^ (dot) chromosome, which has been hypothesized to be an ancestral sex chromosome^14, 15^, while Dyak Caf40 does not. The pattern of bias mirrored that observed for the X chromosome: Dmel Zeus was strongly enriched for signal on the 4^th^ chromosome (p<.001), whereas Dsim Zeus was mildly, but significantly, enriched (p<.05). Both the X and dot chromosomes are enriched for female-biased genes^16^ (Fisher’s Exact Test: p = 2.728×10^-14^), consistent with Zeus’s hypothesized repressive role in the testes^17^. The chromosomal distribution of sites thus suggests a scenario in which Zeus gained affinity for sex-biased genes on the X-chromosome and the 4th chromosome as part of its testis-specific neofunctionalization, and then subsequently evolved differences in chromosome-level binding between *D. melanogaster* and *D. simulans*. Genes on *D. melanogaster’s* 4th chromosome were found to be, on average, more highly female-biased than *D. simulans*, explaining the significant species-specific difference in affinity^18^ (See Supplemental Information; Permutation test, p<.05).

Caf40 is among the most conserved nucleic-acid binding proteins across eukaryotes, from metazoans to fungi to flowering plants^19^. We reasoned that the extensive protein-coding (trans-) divergence of Caf-40 and Zeus may have driven evolution of conserved, bound cis-regulatory elements^20^. We therefore searched for overrepresented motifs for each protein using DREME^21^. A single, highly specific motif (ACTGCTT) was enriched in all three proteins’ binding sites (Supplementary Fig. 5-7). We call this motif the <u>C</u>af-40 <u>a</u>nd <u>Z</u>eus-<u>A</u>ssociated <u>M</u>otif (CAZAM).

We first noted that the genome-wide frequency of the CAZAM differed between *Drosophila* species with and without the Zeus gene. The three species of sequenced Drosophilids with both the *Zeus* and *Caf40* genes had significantly lower overall CAZAM frequencies than sequenced species with only *Caf40*, which remained true after correcting for genome size (Fig. 3; Supplementary Table 2; phylogenetic ANOVA^22^, p=0.004). No randomly constructed motifs were similarly unevenly distributed among the genomes (Supplementary Fig. 8).

**Figure 3.**
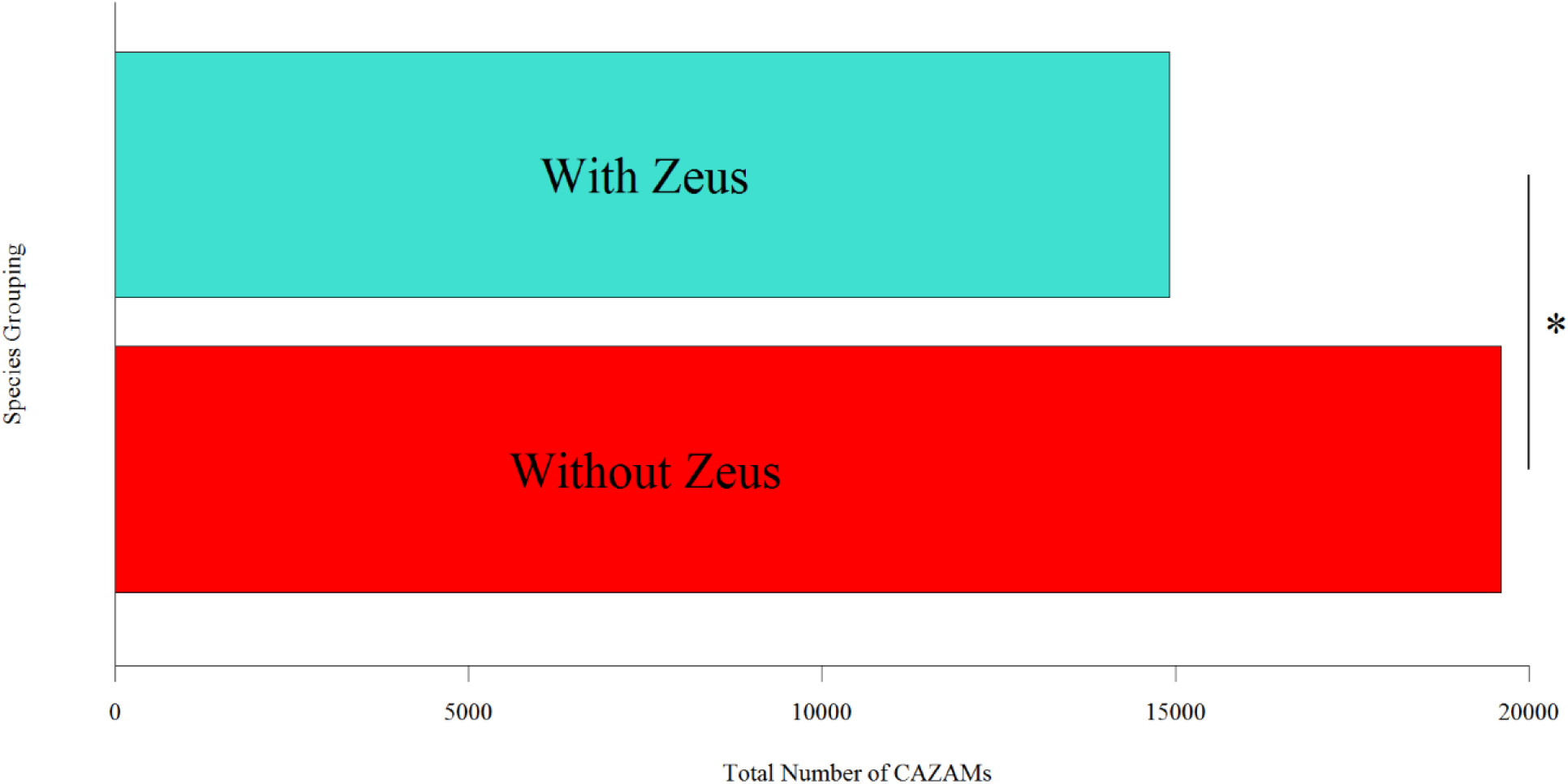
CAZAM Frequency Varies Between Species Which Possess and Do Not Possess Zeus. Bar plot showing the mean frequency of the CAZAM in species with (top bar) and without (bottom bar) the Zeus duplication. The frequency of the CAZAM is significantly lower in species with Zeus (p<.01).

In addition to an overall difference in CAZAM frequency, we found that the distribution of the motif was radically different among the genomes with and without *Zeus*. After the origination of *Zeus*, the frequency of CAZAMs on the X-chromosome did not change appreciably, while motifs decreased on the autosomes (Supplementary Table 2). The fraction of motifs within 1kb of exons, on both X and autosomes, increased dramatically as well (97.8% in *D. simulans* vs. 84.0% in *D. yakuba*; Fisher’s Exact Test: p < 1×10^-7^). The increase in exon-proximal binding may be indicative of a refining of target specificity to those genes required for Zeus’ new function. These results suggest selection acted to remove and redistribute thousands of copies of the motif following the gene duplication event, perhaps as a result of a new regime of selection driven by the emergence of *Zeus*.

To more directly assay for positive selection, we modified the framework of the McDonald-Kreitman test so that it could apply to motif-level analyses at a whole-genome scale^23–26^ (Figure 4A; see Supplemental Information). Because we posited, based on the overall difference in motif frequency between species, that there was selection to remove the motif from the genomes of species after *Zeus* duplicated from *Caf40*, we identified motif instances in preduplication species (*D. yakuba*, *D. pseudoobscura*) as well as one post-duplication species (*D. simulans)* and mapped their syntenic locations into *D. melanogaster*. We note that our version of the test may be conservative, as the flanking regions surrounding motifs showed evidence of stronger purifying selection than synonymous sites, the usual reference for the McDonald-Kreitman test (Akashi 1997).

**Figure 4.**
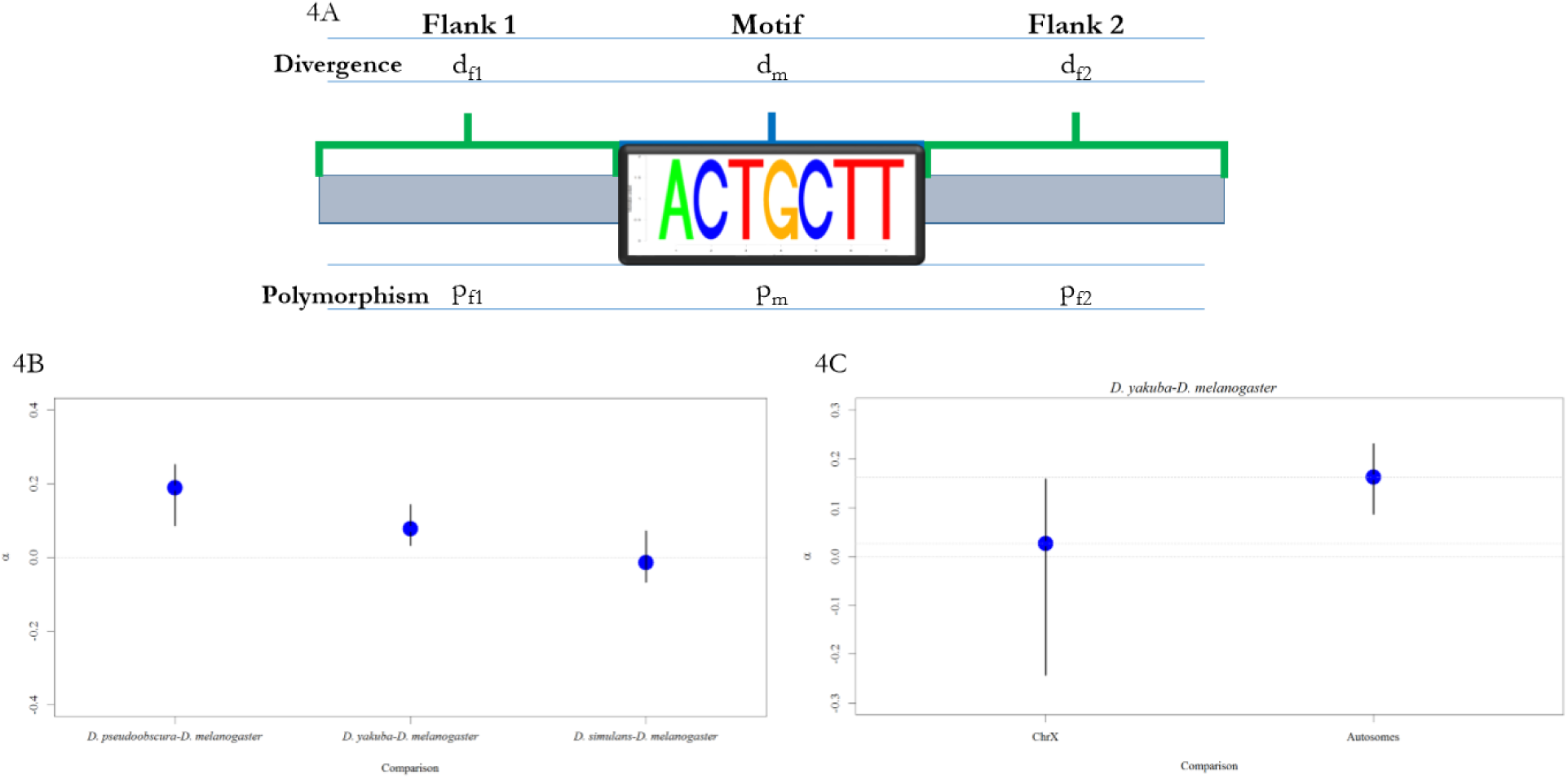
The CAZAM Exhibits a Signature of Selection. A) Schematic illustrating our modification of the McDonald-Kreitman test. We substitute central and flanking sites for dN and ds, respectively, allowing us to measure selection on all identified instances of the motif. B) Plot depicting observed alpha (α) values (interpreted as the proportion of adaptive substitutions) for different comparisons, with bootstrapped 99% confidence intervals. Motifs located and mapped from *D. pseudoobscura* and *D. yakuba* show values of alpha significantly different from zero, while motifs from *D. simulans* do not. C) Comparison of estimated alpha values from motifs which map to the X chromosome (left) in contrast to those which map to the autosomes (right). Values of alpha are significantly lower, indicative of weaker selection, for motifs located on the X chromosome.

Using divergence data from whole-genome multiple alignments between each compared species and *D. melanogaster*, and polymorphism data for *D. melanogaster*^27^, we found that there was significant evidence of positive selection on instances of the CAZAM following the gene duplication event (Figure 2C-D; *D. pseudoobscura-D. melanogaster*. Bootstrap test: p<.01; *D. yakuba-D. melanogaster*. Boostrap test: p<.01). Selection was significantly stronger on intergenic motifs than on exonic motifs, consistent with our findings that all three proteins bound near exonic regions and that there was redistribution of the motif following the duplication event (Supplementary Table 3; p<.01). In contrast, performing the same comparison between *D. simulans* and *D. melanogaster* revealed no significant signature of positive selection, suggesting that strong selection acted after the duplication event, but decreased by the time the *D. melanogaster* and *D. simulans* lineages diverged^3, 28^ (Figure 4B).

Because we determined earlier that Zeus binding shows a strong chromosome-specific bias consonant with its role in testes development^29^, we posited that regimes of selection may have differed across chromosomes. Correspondingly, we found evidence of stronger selection to remove CAZAMs from autosomal chromosomes than from the X chromosome (Fig. 4C; comparing intergenic motifs in *D. yakuba* and *D. melanogaster;* Permutation test: p<.01). We conclude, based on the motif frequency difference and associated evidence of positive selection, that widespread selection driven by the origination of *Zeus* shaped both the abundance and distribution of the motif between species.

To confirm that this signature of selection on the CAZAM was related specifically to the appearance of *Zeus*, we performed a series of control analyses. We examined several motifs that were shuffled versions of the CAZAM, and found significant McDonald-Kreitman test results for none of them after multiple testing correction (Bootstrap tests: p>.5; see Supplemental Information, Supplementary Table **X**). We examined sequence divergence of the CAZAM between two species *(D. pseudoobscura* and *D. yakuba)* without Zeus, and found no significant signal of increased divergence at motif sites relative to flanks (Supplementary Fig. 9; Bootstrap test: p>.5). Our results therefore suggest that it was the origination of *Zeus* that led to widespread positive selection specifically on the CAZAM.

## Discussion

Our results show that Zeus, a novel nucleic-acid binding factor in *Drosophila*, underwent a regime of rapid neofunctionalization ultimately leading to specialized binding to different chromosomes in different species. This trans-evolution in turn drove strong positive selection to rearrange the chromosomal distribution of the motif associated with both Zeus and Caf40 binding. We have thus revealed a dynamic, genome-wide co-evolutionary process of neofunctionalization occurring in both cis and trans.

With regard to the specific molecular mechanism by which Zeus might be regulating downstream targets, initial studies of Caf40 (also known as Rcd-1) suggested that it regulates target genes through direct interaction with the genome due to the fact that it contains six armadillo-type repeats, implicated in DNA binding (Chen et al, 2001; Garces et al, 2007). However, more recent work suggests that Caf40 in Drosophila may also act indirectly with nucleic acids as a member of the larger Ccr4-Not complex, which has roles in mRNA processing and degradation (Bawankar et al, 2013; Collart & Panasenko, 2012; Temme et al, 2010). Our data show that via ChIP, we can indeed discover signals of Caf40 and Zeus binding that illuminate their evolutionary histories, although we cannot discount the possibility that the signals we detect could in fact be due to indirect interactions with the genome mediated through protein-protein binding. For example, extensive studies of transcription factors (TF) binding have revealed that interactions between TFs and the genome are mediated through a highly complex and variable suite of direct and indirect interactions between TFs and cofactors (Slattery et al, 2011; Kazemian et al, 2013; Stefflova et al, 2013; Spitz & Furlong, 2012; Siggers et al, 2011). Although we do not have definitive evidence that Zeus functions within a protein complex, we are confronted by the tantalizing possibility that Zeus has undergone rapid evolutionary changes both in terms of its protein-protein and protein-nucleic acid interactions. Further study of the detailed biochemistry of Zeus should shed light on its precise mechanism of action. We also noted from our analysis that both Dmel and Dsim Zeus show enriched binding on the X and 4th chromosomes, consistent with the putative role of Zeus in the downregulation of female-biased genes. This finding is consonant with the fact that these two chromosomes are heavily hetrochomatinized, and that Zeus may also be involved in chromatin dynamics (Chen REF, Arthur 2014).

Regarding the evolution of the CAZAM, one might suggest that there are several important caveats that apply to our version of the McDonald-Kreitman test. Because of the genome-wide nature of our test, we examined many motifs which are likely not bound by either Caf-40 or Zeus due to occlusion by chromatin or other DNA-bound factors. In addition, extending the M-K test to a genome-wide scale aggregated many unlinked motifs, which can have adverse and unpredictable effects upon the bias of the test (Andolfatto 2008). However, by creating an empirical null distribution of sequences resembling, but different from, the CAZAM, many of the potential issues with the modified M-K test can be reduced. If the test was overly liberal in detecting selection, we would expect to see selection on the permuted CAZAM sequences, as well as in pairs of species which did not differ in terms of the presence of Zeus. Instead, we find that the null hypothesis is rejected only for the specific motif we found in our ChIP-seq data, and only in the particular case in which one compares two species across a specific phylogenetic node which corresponds to the origination of Zeus.

Our results shed light on the fate of newly-arisen functional gene duplicates. From our studies of Zeus, we have demonstrated that novel regulatory proteins may cause positive selection to drive genome-scale rewiring of the transcriptional networks into which they integrate through changes both in the protein itself and the global cis-regulatory environment. These global changes, in turn, can have important phenotypic consequences (e.g. the development and function of the reproductive system), even over relatively short evolutionary time scales.

## Methods Summary

### ChIP-seq Data Production.

ChIP-seq experiments were performed using standard modEncode protocols (www.modencode.org) after collecting adults in each species.

Sequencing data was generated by the High-Throughput Genome Analysis Core (HGAC) at the Institute for Genomics and Systems Biology. All sequencing data is available at GEO, under accession number GSE XXXX-XXX.

### Genomic Data Analysis.

ChIP-seq reads were mapped with BWA^30^, using default parameters, to the most recent UCSC genome versions. Motif discovery was performed with DREME^16^.

### Population Genetics.

We used SNP calls from the DGRP^23^, filtering variants with minor allele frequency less than .05 to remove weakly deleterious variation.

## Acknowledgements.

BHK was supported by an NIH Genetics and Regulation Training Grant (T32GM007197) and a Department of Education GAANN Fellowship (P200A090309/P200A120178). RKA was supported by an NSF Graduate Research Fellowship and an NIH Genetics and Regulation Training Grant (T32GM007197). We thank Steffen Lemke, Al Handler, Lijia Ma, and Daniel Matute for technical advice and for providing experimental materials and reagents. We also thank Rebecca Spokony, Kacy Gordon, Aashish Jha, Sidi Chen, and Grace Yuh Chwen Lee for helpful comments and critical review of drafts of our manuscript. We are grateful to Joe Thornton, Martin Kreitman, and Ilya Ruvinsky for useful advice. We are indebted to innumerable members of the White and Long labs for beneficial criticism, advice, and support.

